# Baseline and Innate Immune Response Characterization of a *Zfp30* Knockout Mouse Strain

**DOI:** 10.1101/2020.06.17.154526

**Authors:** Lucas T. Laudermilk, Adelaide Tovar, Alison K. Homstad, Joseph M. Thomas, Kathryn M. McFadden, Miriya K. Tune, Dale O. Cowley, Jason R. Mock, Folami Ideraabdullah, Samir N. P. Kelada

## Abstract

Airway neutrophilia is correlated with disease severity in a number of chronic and acute pulmonary diseases, and dysregulation of neutrophil chemotaxis can lead to host tissue damage. The gene *Zfp30* was previously identified as a candidate regulator of neutrophil recruitment to the lungs and secretion of CXCL1, a potent neutrophil chemokine, in a genome-wide mapping study using the Collaborative Cross. ZFP30 is a putative transcriptional repressor with a KRAB domain capable of inducing heterochromatin formation. Using a CRISPR-mediated knockout mouse model, we investigated the role that *Zfp30* plays in recruitment of neutrophils to the lung using models of allergic airway disease and acute lung injury. We found that the *Zfp30* null allele did not affect CXCL1 secretion or neutrophil recruitment to the lungs in response to various innate immune stimuli. Intriguingly, despite the lack of neutrophil phenotype, we found there was a significant reduction in the proportion of live *Zfp30* homozygous mutant mice produced from heterozygous matings. This deviation from the expected mendelian inheritance (i.e. transmission ratio distortion) implicates *Zfp30* in fertility or embryonic development. Overall, our results indicate that *Zfp30* is an essential gene but does not influence neutrophilic inflammation in this particular knockout model.

## Introduction

Neutrophils are key participants in the innate immune system’s response to pathogens, but the mechanisms by which these cells respond to immune challenge are prone to generating collateral host tissue damage (Nathan 2006). This makes regulation of neutrophil chemotaxis particularly important in defending against outside insults and preventing unwanted organ damage. This signaling balance is particularly vital in the lungs due to the continual exposure to pathogens, allergens, and other environmental exposures that are introduced through respiration.

Dysregulation of the chemokines that attract neutrophils into tissues may tip the scale from appropriate innate immune response to unwanted damage. The CXC chemokine family members CXCL8 (IL-8), CXCL1 (KC/Gro-α), CXCL2 (MIP-2/Gro-β), and CXCL5 (LIX/ENA-78) are all hallmark neutrophil recruitment molecules that signal through the chemokine receptor CXCR2 (Charo and Ransohoff 2006). Pharmaceutical targeting of CXCR2 has proven effective in reducing airway neutrophilia in early chronic pulmonary disease trials (Kirsten *et al.* 2015; Moss *et al.* 2013; Nair *et al.* 2012; Rennard *et al.* 2015; Todd *et al.* 2016; Watz 2017). CXCR2 antagonism also reduced airway neutrophils in mild atopic asthmatics and patients with severe or persistent neutrophilic asthma (Nair *et al.* 2012; Todd *et al.* 2016; Watz *et al.* 2017).

Previously, we identified the murine gene *Zfp30* as novel regulator of neutrophil recruitment to the airways in a mouse model of asthma (Rutledge *et al.* 2014). Our approach involved use of incipient lines of Collaborative Cross (CC), a multiparental genetics reference population (Srivastava *et al.* 2017), that were treated with the house dust mite allergen Der p 1 (Rutledge *et al.* 2014). In that study, we mapped quantitative trait loci (QTL) to proximal chromosome (Chr) 7 for both neutrophil number and the neutrophil chemokine CXCL in bronchoalveolar lavage fluid. Using expression QTL (eQTL) mapping of whole lung RNA levels, we identified a cis-eQTL for *Zfp30* that co-localized with the neutrophil/CXCL1 QTL. *Zfp30* expression was strongly and negatively corrrelated with neutrophil number and CXCL1 concentration, and in subsequent *in vitro* experiments we showed that decreasing *Zfp30* expression (by siRNA knockdown) lead to increased CXCL1 secretion by airway epithelial cells, consistent with the *in vivo* data. Thus, we concluded that *Zfp30* expression, which we subsequently showed is largely determined by rs51434084 genotype (Laudermilk et al 2018), is a key determinant of CXCL1 and neutrophil recruitment to the airways in response to house dust mite allergen exposure in the CC population. Additionally, because previous studies identified QTL for endotoxin response (Matesic *et al.* 1999) and *Streptococcus pneumoniae* infection (Denny *et al.* 2003) at the same location on Chr 7, we reasoned that ZFP30 may be involved in response to multiple innate immune stimuli.

ZFP30 is a C2H2 zinc finger protein with a KRAB domain. The C2H2 domains allow ZFP30 to bind to DNA in a sequence specific manner, and the KRAB domain recruits KAP1, a well-studied transcriptional repressor to these binding sites (Friedman *et al.* 1996). This transcriptional repression proceeds through recruitment of HP1, SETDB1, and histone deacetylases that induce heterochromatin formation and silence nearby genes (Groner *et al.* 2010; Medugno *et al.* 2005; Ryan *et al.* 1999; Schultz *et al.* 2002;). There is precedent for zinc finger proteins to regulate immune signaling through heterochromatin domains, including ZNF160 which down-regualtes TLR4 expression in intestinal epithelia to ameliorate immune response to the host microbiome (Takahashi *et al.* 2009).

In this study, we further examined *Zfp30* and its role in neutrophil recruitment to the airways through generation of a CRISPR-Cas9 *Zfp30* knockout mouse strain. Given that we previously found that low *Zfp30* expression was associated with higher neutrophil counts in the airways after exposure to house dust mite allergen, we hypothesized that *Zfp30* knockout mice would exhibit heightened levels of neutrophilic inflammation after treatment with allergen and other innate immune stimuli. Additionally, because a recent report described a regulatory effect of ZFP30 on *Pparg2* expression and adipogenesis (Chen 2019), we also assessed metabolic phenotypes in *Zfp30* wildtype and knockout mice. Somewhat surprisingly, we found that the CRISPR-Cas9 *Zfp30* null allele resulted in non-mendelian inheritance, specifically transmission ratio distortion, with significant depletion of homozygous *Zfp30* knockout mice in matings of *Zfp30^+/-^* x *Zfp30*^+/-^ mice. However, contrary to expectation, we found that *Zfp30* knockout mice did not exhibit differences in neutrophil recruitment to the airways after exposure to innate immune stimuli, nor did we observe any differences in metabolic phenotypes or *Pparg2* expression in adipose tissue from *Zfp30* knockout mice.

## Materials and Methods

### Generation of *Zfp30* Knockout Mice by CRISPR/Cas9 Embryo Microinjection

A CRISPR/Cas9 guide RNA (5’-GAATCCAGATACAGCAGTAA(CGG)-3’) was designed to target mouse (NCBI Taxon ID: 10090) *Zfp30* gene (NCBI Gene ID: 22693) near the 5’ end of exon 5. While targeting earlier exons would have been preferable, exon 5 was the only region that could be targeted with specificity owing to high homology across ZFP family members. Exon 5 encodes the C2H2 zinc finger domains of ZFP30 that are required for DNA binding. The guide RNA was produced by T7 *in vitro* transcription and validated *in vitro* by incubating guide RNA, Cas9 enzyme, and plasmid harboring the guide RNA target site. This was followed by gel electrophoresis to determine the extent of in vitro cleavage activity. A donor oligonucleotide (“Zfp30-H1-T”: 5’-GTTTTTCTTCTTTTTGCTTTCAGATCTGGAATCCAGATACAGC[TGA][TA<u>G]GATC</u> <u>C</u>[TAG]ACCGGTAACGGGTTACTTCCAGAAAAGAATACTTACGAAATTAATCTATCT-3’) was used for homologous recombination to insert stop codons (brackets) and a BamHI restriction site (underlined) at the target site (Supplementary Figure 1).

C57BL/6J females were then superovulated by injection with pregnant mare’s serum gonadotropin (PMSG) and human chorionic gonadotropin and then mated with C57BL/6J stud males for zygote production. One-cell embryos were collected from the ampulla oviducts the morning after mating and microinjected with either “low” or “high” mix, containing, respectively, 20 ng/μl or 100 ng/μl in vitro transcribed Cas9 mRNA, 20 ng/μl or 50 ng/μl Zfp30 guide RNA and 100 ng/μl Zfp30-H1-T donor oligonucleotide. The microinjected embryos were then implanted into pseudopregnant recipients.

Fourteen live pups born from microinjected embryos were screened by polymerase chain reaction (PCR) amplification of the *Zfp30* target site followed by digestion of the PCR product with BamHI restriction enzyme. The BamHI restriction site was detected in nine animals. Two founders with apparent biallelic insertion of the BamHI restriction site were mated to C57BL/6J animals for germline transmission of the targeted allele.

The founder animals harboring the intended *Zfp30* mutant allele were screened for mutations at 10 potential off-target sites (Supplementary Table 1). Each potential off-target site was PCR amplified and products were analyzed by T7endo1 assay. Founders chosen for line establishment were further analyzed by Sanger sequencing of PCR products for all 10 off-target sites. A single founder line was subsequently backcrossed to C57BL/6J again to remove detected off-target mutations. The mutant strain, C57BL/6J-*Zfp30^em1Snpk^*/Mmnc (hereafter referred to simply as *Zfp30*^-/-^), has been deposited into the Mutant Mouse Research and Resource Center at UNC (https://www.mmrrc.org/catalog/sds.php?mmrrc_id=50629).

In the initial stages of breeding, genotyping of *Zfp30*^-/-^ mice was performed using allele-specific PCR. Primer sets were designed to specifically target either the *Zfp30*^+/+^ (Fwd: GGGCTGCTAAGTCCATTCAG; Rev: GGAAGTAACCCGTTACTGCTG) or *Zfp30*^-/-^ (Fwd: GGGCTGCTAAGTCCATTCAG; Rev: CGTTACCGGTCTAGGATCCT) allele. We later transitioned to a proprietary qPCR-based genotyping protocol through Transnetyx (Cordova, TN).

### qPCR for *Zfp30* Quantification in the Knockout Strain

To quantify *Zfp30* gene expression level in *Zfp30*^+/+^ and *Zfp30*^-/-^ mice, we designed primer sets that specifically quantify the *Zfp30*^+/+^ allele (Fwd: TGTTGGAACAAGGGAAGGAG; Rev: GTAACCCGTTACTGCTGTAT) or specifically quantify the *Zfp30*^-/-^ allele (Fwd: TGTTGGAACAAGGGAAGGAG; Rev: CGGTCTAGGATCCTATCAGCT). qPCR reactions were carried out using iTaq Universal SYBR Green Supermix (Bio-Rad; Hercules, CA USA).

### Complete Blood Count Assays

For complete blood counts, blood was collected in EDTA tubes and stored on ice for a minimal amount of time before processing via a ProCyte Dx Hematology Analyzer.

### Metabolic phenotyping

Magnetic Resonance Imaging (MRI)I: MRIs scans were performed using an EchoMRI-3n1-100TM analyzer prior to glucose tolerance tests to accurately determine body mass composition.

Glucose Tolerance Test: At 13 weeks of age, mice were fasted overnight and dosed with glucose (2g per kg lean body mass) via intraperitoneal injection. Blood glucose was measured at baseline and 15, 30, 45, 60, and 120 minutes after injections, using an Accu-Chek Performa glucometer and test strips (Roche, Basel Switzerland).

### Neutrophil Recruitment Models

Lipopolysaccharide (LPS) challenge: Intratracheal instillation of LPS from *E. coli* (LIST Biologicals Campbell, CA) into lungs of *Zfp30*^+/+^ and *Zfp30*^-/-^ mice was carried out at a dose of 0.3 mg per kg of body weight using previously described methods (Limjunyawong *et al*. 2015, Mock *et al*. 2020). Bronchoalveolar lavage fluid was collected between 8 and 48 hours after exposure, and differential cell counts in bronchoalveolar lavage fluid were performed. Aliquots of BALF were saved for cytokine quantification. Oropharyngeal aspiration of LPS from *E. coli* into lungs of *Zfp30*^+/+^ and *Zfp30*^-/-^ mice was carried out with 5μg LPS in 40μl PBS.

*Dermatophagoides pteronyssinus* house dust mite allergen (Der p 1): We used a model of allergic inflammation involving Der p 1 that we previously showed induces predominantly eosinophilic but also neutrophilic inflammation (Kelada et al. 2011). *Zfp30*^+/+^ and *Zfp30*^-/-^ mice were sensitized with 10 μg Der p 1 (Indoor Biotechnologies, Charlottesville, VA) administered through intraperitoneal injection (in 100 μl of PBS) on days 0 and 7 of the experiment, and a 50 μg Der p 1 challenge was administered on day 15 of the experiment (Kelada *et al.* 2011). Mice were sacrificed 48-72 hours after challenge, and differential cell counts in bronchoalveolar lavage fluid were performed. Aliquots of BALF were saved for cytokine quantification.

Ozone exposure: *Zfp30*^+/+^ and *Zfp30*^-/-^ mice were exposed to filtered air, 1 ppm ozone, or 2 ppm ozone for three hours as previously described (Smith *et al.* 2019). Bronchoalveolar lavage fluid was collected 24 hours after exposure, and differential cell counts in bronchoalveolar lavage fluid were performed.

### Mouse tracheal epithelial cell (MTEC) culture model

MTEC cultures were generated and cultured according to a previously established protocol (You and Brody 2013). Tissues isolated from 4 week old male and female mice were grown using PluriQ differentiation media and plated in 12 well plates with Transwell inserts. Cells were maintained at air-liquid interface for a minimum of three weeks to allow for differentiation. LPS, a TLR4 ligand that induces strong CXCL1 secretion, exposures were carried out at 10μg/mL in 100μl of PBS added to the apical surface of MTECs for 24h.

### Luminex Assays

Cytokines in bronchoalveolar lavage fluid or PBS used in MTEC LPS exposures were measured using Milliplex assays (Millipore, Billerica, MA) according to manufacturer’s instructions.

### Histology

Histological preparation and analysis of lungs were carried out using previously described methods (Donoghue *et al.* 2017). Briefly, left lung lobes were fixed in formalin and cut in cross-section starting at the hilum and 2mm apart along the main stem bronchus. Sections were embedded in paraffin and stained with Hematoxylin and eosin (H&E) stain. Images were captured on an Olympus BX605F microscope with CellSens Standard software.

## Results

We sought to investigate the function of *Zfp30* using a CRISPR-Cas9 generated *in vivo* knockout model. Of fourteen pups born from microinjected embryos, nine had successful insertion of the donor oligonucleotide harboring the BamHI restriction site, and two founders with biallelic insertion were mated to C57BL/6J for germline transmission of the targeted allele. These founder animals were screened for ten potential off-target mutations. Bi-allelic insertion/deletion (indel) mutations were identified at a single off-target site in both founders. N1 backcross animals from a single founder were subsequently backcrossed once more to C57BL/6J to remove the off-target indel mutation and establish our *Zfp30*^+/-^ colony.

Throughout the maintenance of this colony, through breeding of heterozygous animals, transmission ratio distortion (TRD) was a recurring issue. A substantial depletion of homozygous knockout animals was first apparent in the early stages of breeding, and this depletion was initially overcome through selective breeding of mice that produced litters of normal genotype distribution. However, TRD appeared again after several generations of breeding. Among a total of 1,270 mice produced from *Zfp30*^+/-^ x *Zfp30*^+/-^ matings, only 289 (22.8%) offspring were homozygous knockouts (Table 1). Using the binomial distribution formula with genotype frequency expectations based on mendelian inheritance (i.e., 25% frequency of homozygous knockout mice), we calculated the cumulative probability of observing 289 or fewer homozygous mutant mice to be 0.03. Further examination revealed that this TRD appeared to solely impact female pups (Table 1). Of 648 female mice produced from *Zfp30*^+/-^ x *Zfp30*^+/-^ matings, only 21% were homozygous knockout genotypes (p=0.01 by binomial distribution test). It is unclear if *Zfp30*^-/-^ animals died during embryonic development or if alleles are selected against through some other mechanism. However, mean litter size for the colony is 7.15, and we did not observe any evidence that *Zfp30*^-/-^ mice died after birth. Recently, the International Mouse Phenotyping Consortium generated a *Zfp30* knockout strain and also found that among litters from heterozygous parents there was a significant depletion of homozygous knockout mice (www.mousephenotype.org, Dickinson *et al.* 2016), providing a replication of our result.

**Table 1.** Genotype counts and ratios of all mice born from heterozygous matings

|  | *Zfp30*<sup>+/+</sup> | *Zfp30*<sup>+/-</sup> | *Zfp30*<sup>-/-</sup> | **Total** |
| --- | --- | --- | --- | --- |
| **Male** | 152<br>(24.44%) | 318<br>(51.13%) | 152<br>(24.44%) | 622 |
| **Female** | 185<br>(28.55%) | 326<br>(50.31%) | 137<br>(21.14%) | 648 |
| **Total** | 337 | 644 | 289 | 1270 |

Because no reliable antibody for mouse ZFP30 exists, we verified the *Zfp30* knockout at the RNA level in whole lung tissue from 9-10 week old mice. We designed a qPCR-based approach to specifically quantify expression of either the *Zfp30* wildtype allele or the CRISPR-Cas9 modified *Zfp30* mutant allele. We detected expression of only the *Zfp30* wildtype allele in WT mice, only the *Zfp30* mutant allele in KO mice, and intermediate expression of the two alleles in heterozygous mice (Figure 1).

**Fig. 1.**
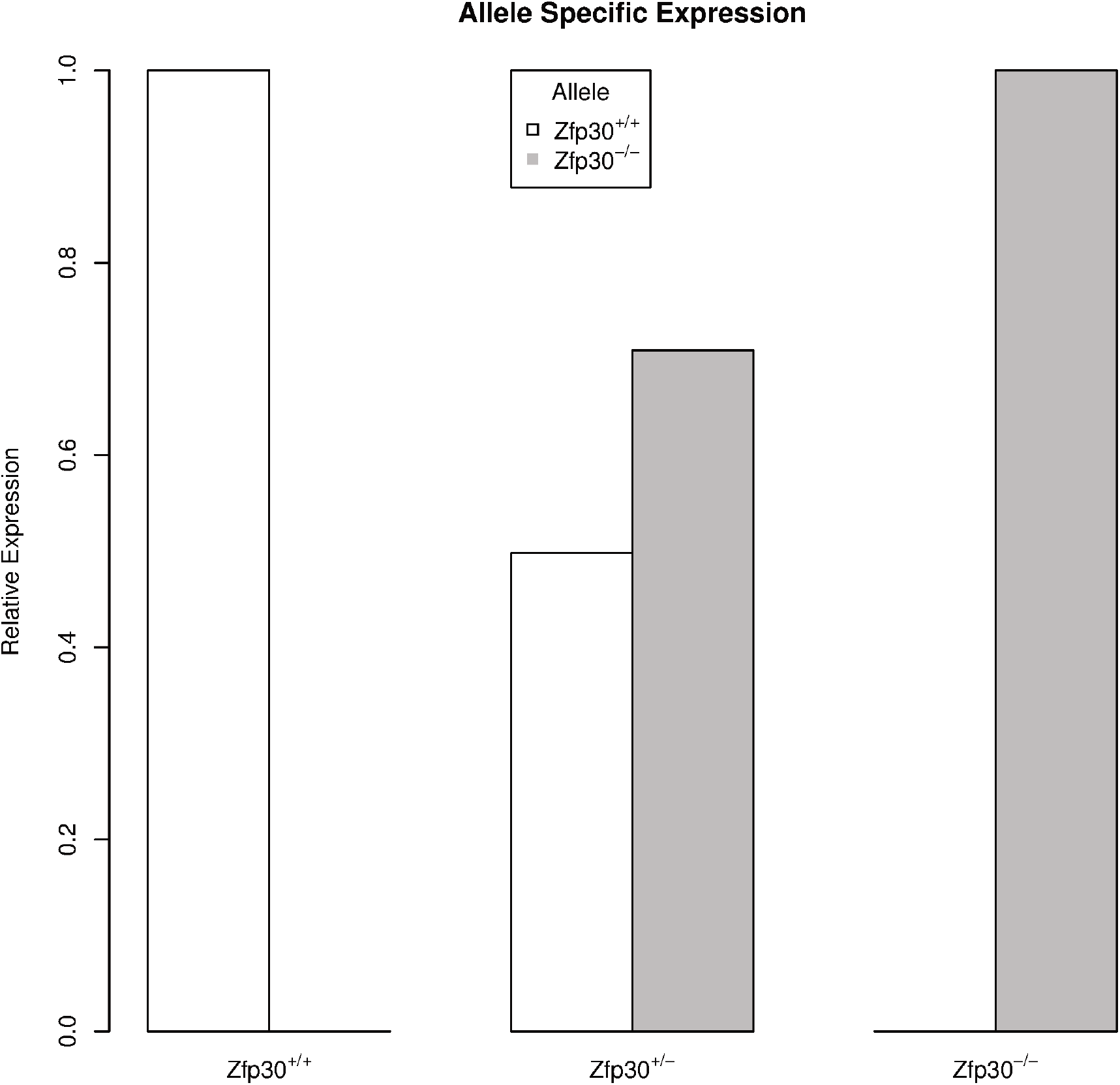
Allele-specific qPCR confirms genotype-dependent expression of *Zfp30* wildtype and mutant alleles in the lung. Primer pairs that specifically amplify either the wildtype or CRISPR-Cas9 modified allele were used to confirm the status of *Zfp30* expression in whole lung tissue from 4-6 mice per genotype. Mice were 9-10 weeks of age at harvest.

### Baseline Phenotyping

As a family, the KRAB domain-containing C2H2 zinc finger proteins are thought to play an important role in differentiation and development (Lupo *et al.* 2013; Urrutia 2003). To assess the impact that a whole-body knockout of *Zfp30* might have on development, we assayed the weights of lungs, pancreases, spleens, and livers in *Zfp30*^+/+^ and *Zfp30*^-/-^ mice, and we detected no significant differences in these phenotypes (Supplementary Table 2). Complete blood count (CBC) assays did not reveal any significant differences in red blood cell or circulating leukocyte phenotypes between *Zfp30*^+/+^ and *Zfp30*^-/-^ samples (Supplementary Table 3). Because ZFP30 has been reported to play a role in adipogenesis via regulation of *Pparg2* expression (Chen 2019), we tested for metabolic impacts of *Zfp30* knockout in fasting glucose and glucose tolerance (Figure 2A) but saw no significant differences. We did, however, observe a significant difference in the body weights of *Zfp30*^+/+^ and *Zfp30*^-/-^ male mice (Figure 2B) and a marginally significant difference in the lean body weights among male mice (Figure 2C). We did not, however, observe a substantial impact of *Zfp30* knockout on fat mass (Figure 2D) or *Pparg2* expression in white adipose tissue (Fold change (KO vs WT)= 0.82; p-value = 0.62; WT n=11, KO n=10).

**Fig. 2.**
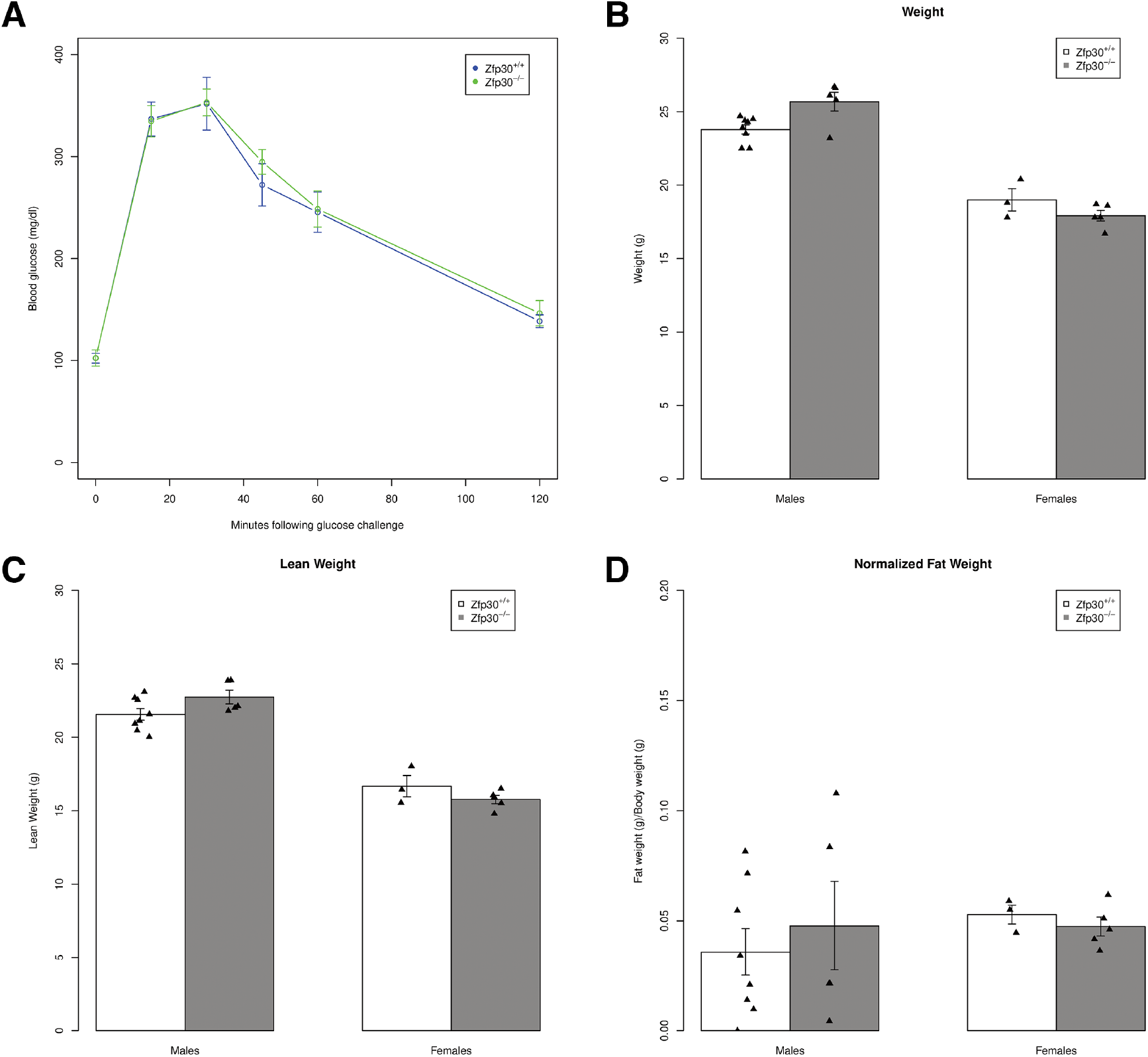
*Zfp30*^-/-^ mice exhibit no impact on glucose tolerance and have lower body weights. **A.** Thirteen week old mice were fasted overnight and dosed with glucose (2 g per kg lean body mass) via intraperitoneal injection. Blood glucose was then measured at baseline and 15, 30, 45, 60, and 120 minutes after injections. **B.** Bodyweight at harvest. **C.** Lean body mass determined by MRI. **D.** Fat mass determined by MRI normalized to body weight. N=10-11 per genotype. * *P* < 0.05; *P* < 0.1.

Finally, we carried out histological analysis of lungs from *Zfp30*^+/+^ and *Zfp30*^-/-^ lungs to investigate any obvious differences in the airways, alveoli, or vasculature and detected no striking differences (Supplementary Figure 2).

### *Ex vivo* Mouse Tracheal Epithelial Cell Cultures

Given our previous results indicating a correlation between *Zfp30* expression and innate immune response in the lung (Rutledge *et al.* 2014), we tested innate immune responses in mouse tracheal epithelial cultures (MTECs) from *Zfp30*^+/+^ and *Zfp30*^-/-^ mice. This system was particularly well suited to study the impact of *Zfp30* knockout on neutrophil recruitment, because recent single-cell RNA sequencing (RNA-seq) data suggests expression of *Zfp30* across a broad array of cell types in the airway epithelium (Plasschaert *et al.* 2018). Additionally, we previously showed that MTEC cultures have high expression of *Zfp30* and that perturbation of *Zfp30* expression with siRNAs results in increased CXCL1 production following LPS exposure in a mouse airway epithelial cell line (Rutledge *et al.* 2014). After establishing MTEC cultures, we tested whether there were differences in the proportions of airway epithelial cell types using qRT-PCR for markers of ciliated cells (*Foxj1*), club cells (*Scgb1a1*), goblet cells (*Muc5ac*), and basal cells (*Krt5*). We detected a marginally significant doubling of *Muc5ac* expression in *Zfp30*^-/-^ MTECs (Table 2), suggesting elevated goblet cell numbers in the knockout. We then stimulated *Zfp30*^+/+^ and *Zfp30*^-/-^ MTECs with LPS, but did not detect significant differences in CXCL1 secretion in response (Figure 3).

**Fig. 3.**
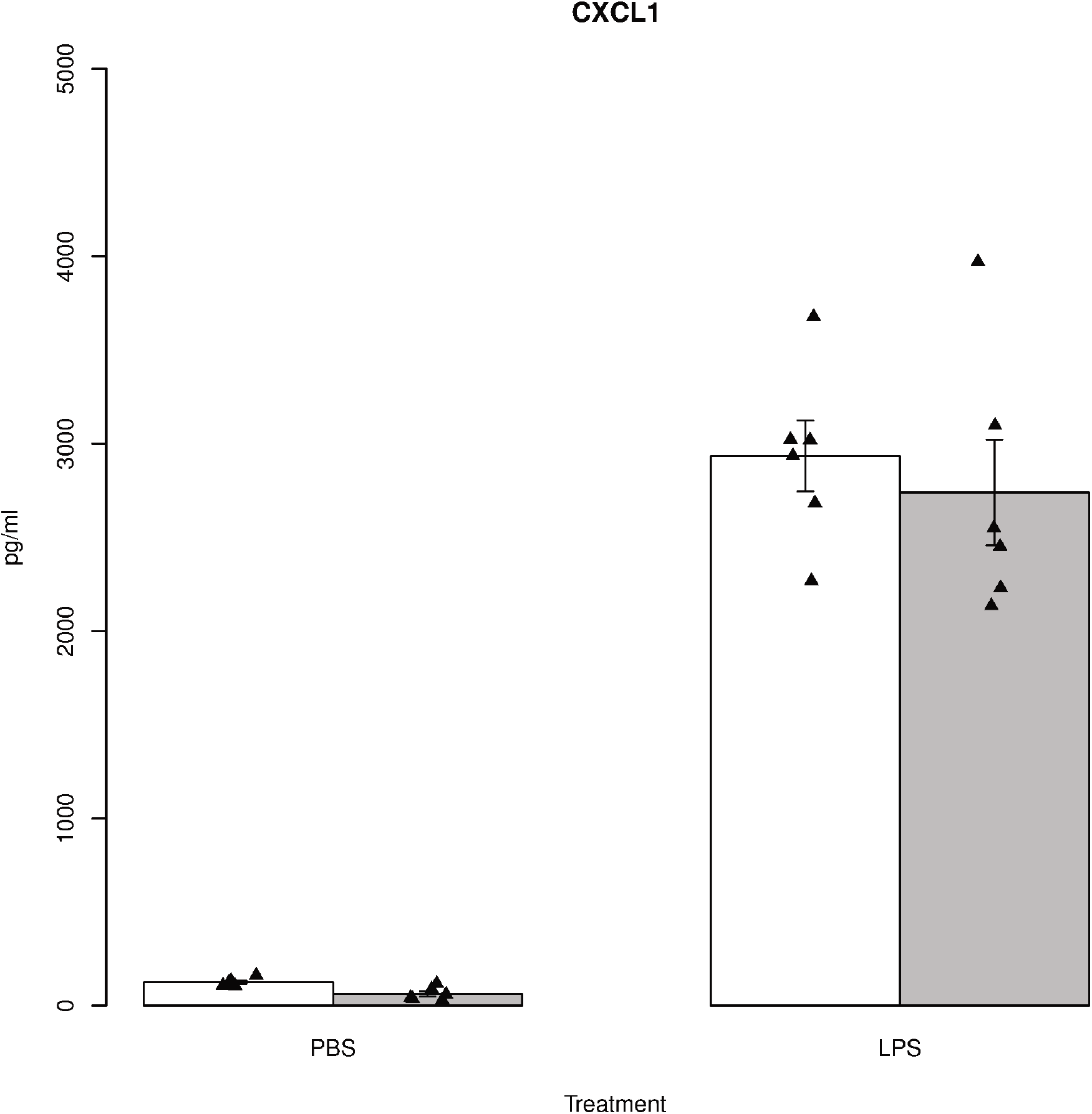
No difference in CXCL1 secretion by genotype in primary mouse tracheal epithelial cells after LPS exposure. Mouse tracheal epithelial cell cultures were generated from *Zfp30*^+/+^ and *Zfp30*^-/-^ mice and maintained at air-liquid interface for 21 days. Cells were exposed 100 μM LPS for 24 hours, then supernatants were assayed for CXCL1 concentrations using Luminex assays. N= 6 per condition per genotype.

**Table 2.** Airway epithelial cell marker gene expression in primary tracheal epithelial cell cultures from *Zfp30*^-/-^ vs. *Zfp30*^+/+^ mice.

| **Sample** | *Foxj1* | *Krt5* | *Scgb1a1* | *Muc5ac* | *Tgm2* | *Lyz2* | *Lif* |
| --- | --- | --- | --- | --- | --- | --- | --- |
| Fold Change* | 0.95 | 1.24 | 1.29 | 2.09 | 1.18 | 1.12 | 0.82 |
| p-value | 0.48 | 0.48 | 0.44 | 0.07 | 0.25 | 0.76 | 0.28 |

\* Fold change for *Zfp30*<sup>-/-</sup> vs. *Zfp30*<sup>+/+</sup> mice using *Abhd6* for normalization.

### *In vivo* Lung Inflammation Phenotypes in *Zfp30*^-/-^ mice

*Zfp30* was identified as a candidate regulator of CXCL1 levels and neutrophils in bronchoalveolar lavage fluid in the context of a model of allergic airway disease (Rutledge *et al.* 2014). As a direct follow-up to these experiments, we utilized the same house dust mite model of allergic airway disease in *Zfp30*^+/+^ and *Zfp30*^-/-^ mice to further probe the connection between *Zfp30* and neutrophil recruitment. We saw no significant differences in neutrophil counts or CXCL1 in bronchoalveolar lavage fluid (BALF) 48 or 72 hours post-challenge (Figure 4A-B).

**Fig. 4.**
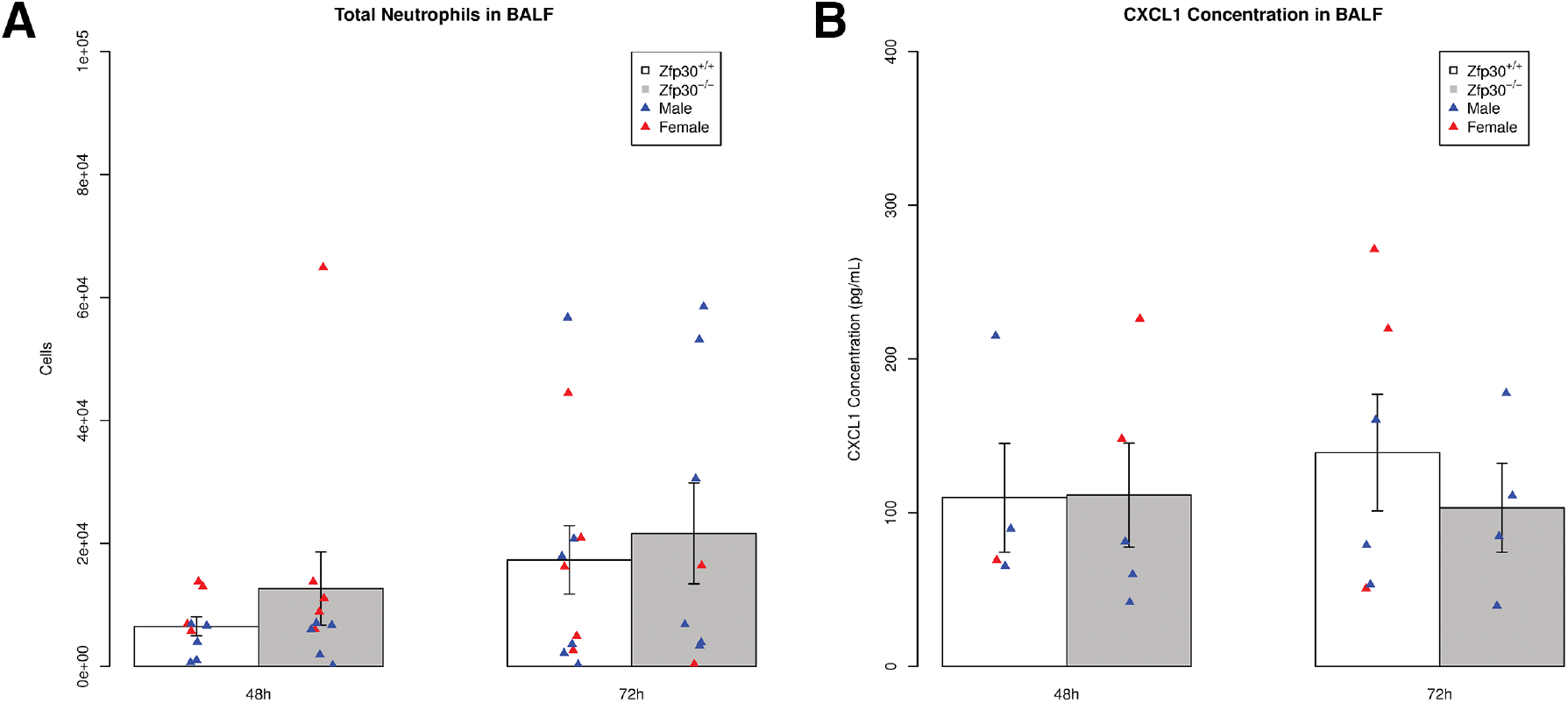
No difference in neutrophil count or CXCL1 concentration in BALF after house dust mite allergen challenge in *Zfp30*^+/+^ and *Zfp30*^-/-^ mice. *Zfp30*^+/+^ and *Zfp30*^-/-^ mice were sensitized to and challenged with HDM allergen, and effects on neutrophil counts (A) and CXCL1 concentration (B) in BALF were monitored 48 and 72 hours after challenge. Cell count results are the combined data of two experiments and were assayed via differential cell counts. N= 9-11 mice per condition per genotype. Cytokine measurements were carried out via multiplex assays. N= 9-10 mice per genotype.

The allergic airway disease model we employed is dominated by eosinophilia. To test for differences in CXCL1 secretion or neutrophil recruitment into the lungs in the context of neutrophil-dominated immune responses, we employed LPS and ozone exposure models. These models induce a much more robust neutrophilic airway recruitment than the HDM model, so differences in chemotactic signaling may be more apparent. We found that intratracheal instillation of LPS did not cause a significant difference in neutrophil or CXCL1 levels in BALF of *Zfp30*^-/-^ vs. *Zfp30*^+/+^ mice (Figure 5A-C). Additionally, there were no significant differences in neutrophilia by genotype in a model of sterile inflammation induced by the air pollutant ozone (1 and 2 parts per million concentration).

**Fig. 5.**
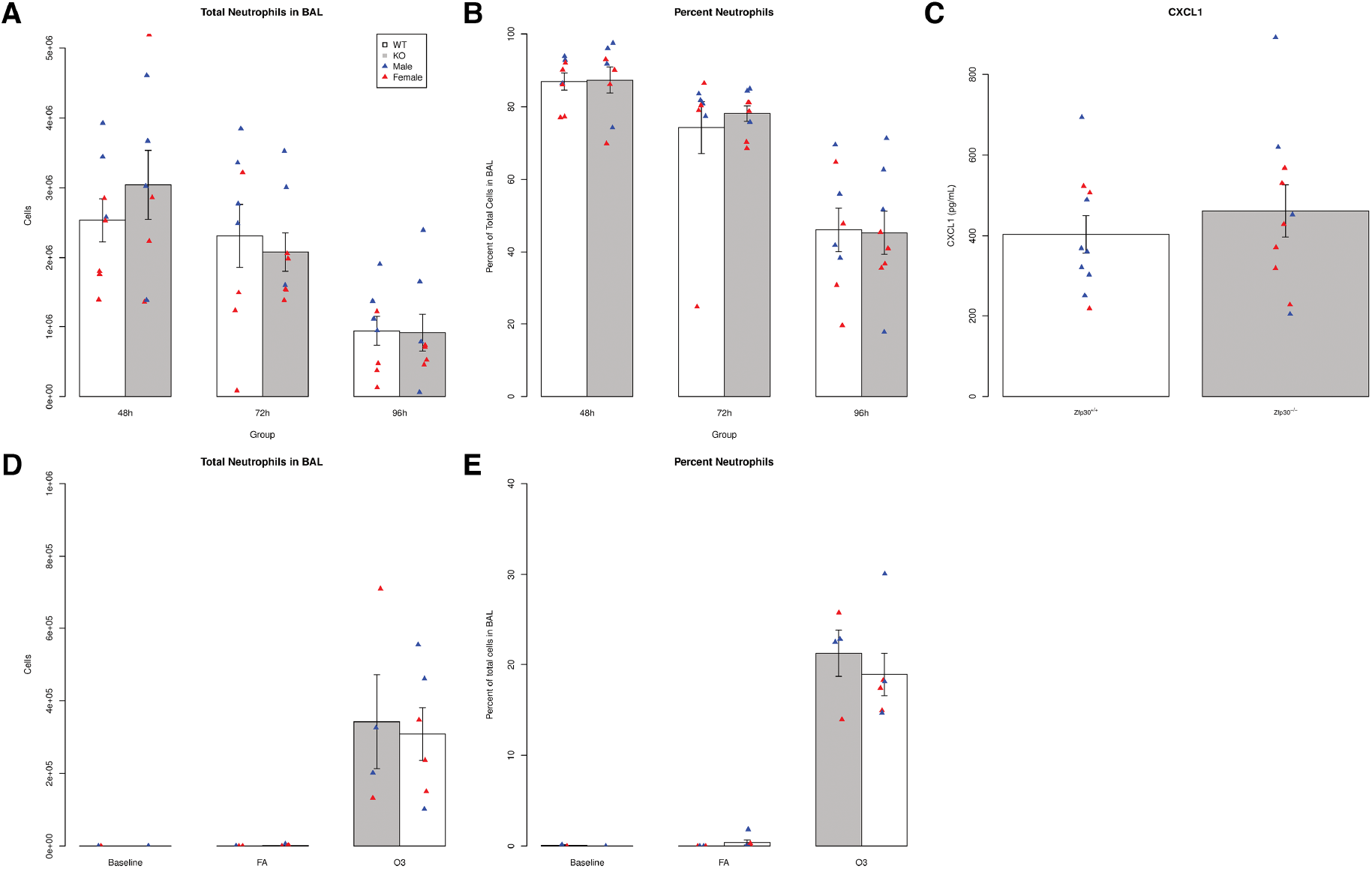
No differences in neutrophil chemotaxis by *Zfp30* genotype in two models of neutrophilic inflammation. *Zfp30*^+/+^ and *Zfp30*^-/-^ mice were given LPS by intratracheal administration and phenotyped 48, 72, or 96 hours later (A-C). Neutrophils counts (A) and neutrophils as a percentage of total cell count in bronchoalveolar lavage (B) were we measured at indicated time points. CXCL1 was measured at 48 hours post-challenge (C). Neutrophil counts reported are from a single intratracheal administration and are representative of four experiments that span collections 8-96 hours after challenge. Neutrophil levels were assayed via differential cell counts. N= 8 per genotype per condition. Cytokine measurements were carried out via Luminex assays. N= 10 per genotype. *Zfp30*^+/+^ and *Zfp30*^-/-^ mice were exposed to 2 ppm ozone for three hours and phenotyped 21 hours later. Total (D) and percent neutrophils (E) are shown. N= 4-6/genotype/group.

## Discussion

Based on previous data that implicates *Zfp30* in the regulation of neutrophil chemotaxis, we developed a *Zfp30* knockout mouse strain to test for potential impacts on *in vivo* neutrophil recruitment. We accomplished this through CRISPR-Cas9 targeting of the DNA binding domains within *Zfp30*. This strategy was chosen to disrupt wiltype DNA binding activity of ZFP30, a crucial component of this transcription factor family’s sequence-specific chromatin remodeling activity. Though no suitable antibody for ZFP30 exists, allele-specific qPCR revealed a total loss of wildtype allele expression in homozygous mutant mice. Contrary to expectation based on our previous work (Rutledge 2014), the results shown here indicate that loss of wiltype ZFP30 function does not significantly affect CXCL1 secretion or neutrophil chemotaxis in response to various innate immune stimuli.

One potential explanation for why ZFP30 loss did not have the predicted effects on neutrophil chemotaxis is that genetic background, i.e., strain, could have altered the impact of *Zfp30* genotype. We generated the *Zfp30* knockout on the C57BL/6J genome, which is one of the Collaborative Cross founder strains. This inbred strain, however, was recently shown to harbor a mutation in *Nlrp12* that impacts neutrophil recruitment (Hornick 2017; Ulland *et al.* 2016). Introducing a knockout into an already-mutated cytokine secretion pathway may have masked the effects of the knockout. Hence, it is possible that generating a *Zfp30* knockout on another genetic background could reveal different phenotypic consequences, as has been observed in other cases (Sittig et al. 2016). *Zfp30* is more highly expressed in the lungs of 129S1/SvImJ, A/J, NOD/ShiLtJ, and NZO/HILtJ laboratory mouse strains, so they may be good candidates for further analysis. Since we have generated a full-body, non-conditional knockout, it is also possible that in *Zfp30*^-/-^ mice, other genes compensated for *Zfp30*, thereby masking any effects of ZFP30 absence (Rossi 2015).

Our analysis of MTEC cultures revealed a marginally significant difference in expression of *Muc5ac* among *Zfp30*^-/-^ cultures, suggesting a possible difference in goblet cell abundance between *Zfp30*^+/+^ and *Zfp30*^-/-^ mice. Given the important role of MUC5AC is mucus hypersecretion and airway obstruction (Evans *et al*. 2015, Ordoñez *et al.* 2001), this finding may merit further investigation. For example, determining whether loss of ZFP30 affects airway epithelial progenitor cell differentiation, or whether *Muc5ac* expression was increased due to some inflammatory process in these cells.

*Zfp30* was recently shown to affect adipogenesis and *Pparg2* expression *in vitro* (Chen 2019), and human *ZFP30* is differentially expressed in the pancreatic beta cells of type-2 diabetes patients (Lawlor 2017). Additionally, previous results suggest that ZFP148 regulates *Zfp30* expression (Laudermilk et al. 2018), and *Zfp148* was recently implicated in glucose tolerance and insulin secretion from pancreatic islets in mice (Keller et al. 2019). However, follow-up studies on insulin sensitivity here revealed no metabolic impact of ZFP30 loss, nor did we observe a significant difference in *Pparg2* expression in the white adipose tissues of our mice. We did, however, observe differences in body weight and lean body weight among male *Zfp30*^-/-^ mice. In contrast to our results, male *Zfp30* knockout mice generated by International Mouse Phenotyping Consortium exhibited decreased fasting glucose concentrations compared to wildtype mice.

One of the most intriguing findings generated here was that there was a significant depletion of homozygous knockout offspring from *Zfp30*^+/-^ x *Zfp30*^+/-^ matings, a finding that was independently reproduced by the International Mouse Phenotyping Consortium. These results implicate ZFP30 in fertility or embryonic development and demonstrate that *Zfp30* is an essential gene. The cause of the transmission ratio distortion and its sex-specific effect remains unclear, especially in light of the fact that litter sizes were not obviously affected. That said, there is precedent for similar effects of other zinc finger protein knockout mice to display some degree of embryonic lethality. Knockout of *Zfp57*, a C2H2 ZFP with a KRAB domain, affects the establishment and maintenance of critical DNA methylation imprints, and disruption of this imprinting leads to death (Li *et al.* 2008). Further studies will be required to explore the mechanisms underlying this phenotype.

To conclude, we demonstrate here that a knockout of *Zfp30* on a C57BL/6J genetic background does not affect CXCL1 secretion or neutrophil recruitment to the lungs in mouse models of asthma or acute lung injury. *Zfp30*^-/-^ male mice do differ from their wildtype littermates in body weight but not in baseline metabolic phenotypes assayed here. Finally, the *Zfp30* null allele is associated with transmission ratio distortion, with a significant depletion of *Zfp30*^-/-^ mice from *Zfp30*^+/-^ x *Zfp30*^+/-^ matings, though the mechanism underlying this depletion remains to be determined.

## Acknowledgments

The authors would like to thank: Gregory J. Smith, Ph.D. for his assistance with *in vivo* ozone exposures; Larry Ostrowski, Ph.D. and Ximena Bustamante, Ph.D. for their assistance with MTEC isolation and culture; Kim Burns for her assistance with histology; Max Lowman for technical assistance with qPCR work; Autumn Sanson for her assistance with *in vivo* data processing; Gang Chen for his insights into mouse tracheal epithelial cell qPCR; and Praveen Sethupathy, Ph.D. and Yu-Han Hung, Ph.D. for their consultation on metabolic phenotypes. The authors would additionally like to thank the UNC CGIBD Advanced Analytics core for their work on cytokine multiplex assays, the Animal Histopathology and Laboratory Medicine core for their work in processing complete blood count assays,and the UNC NORC Animal Metabolism Phenotyping core for their work on mouse MRIs.

## Declarations

### Ethics approvals

Not applicable

### Consent to participate

Not applicable

### Consent for publication

Not applicable

### Availability of data and material

The datasets generated during and/or analysed during the current study are available from the corresponding author on reasonable request.

### Code availability

Not applicable

### Author contributions

All authors contributed to the study conception and design. Material preparation, data collection and analysis were performed by Lucas T. Laudermilk, Adelaide Tovar, Alison K. Homstad, Joseph M. Thomas, Kathryn M. McFadden, Miriya K. Tune, Dale O. Cowley, Jason R. Mock, and Samir N. P. Kelada.

---

**Supplementary Table 1.**
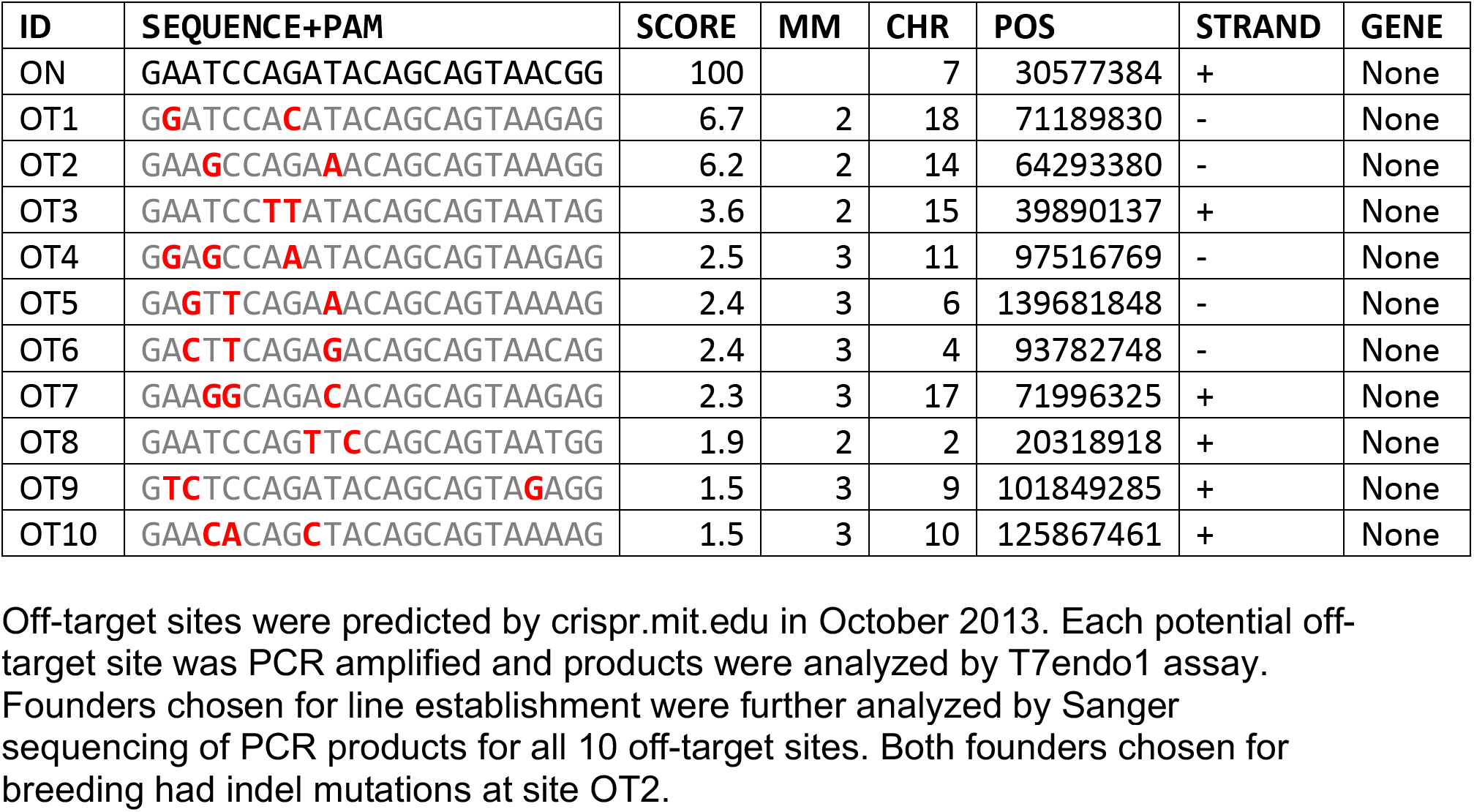
The founder animals harboring Zfp30 mutations were screened for mutations at 10 potential off-target sites

**Supplementary Table 2:**
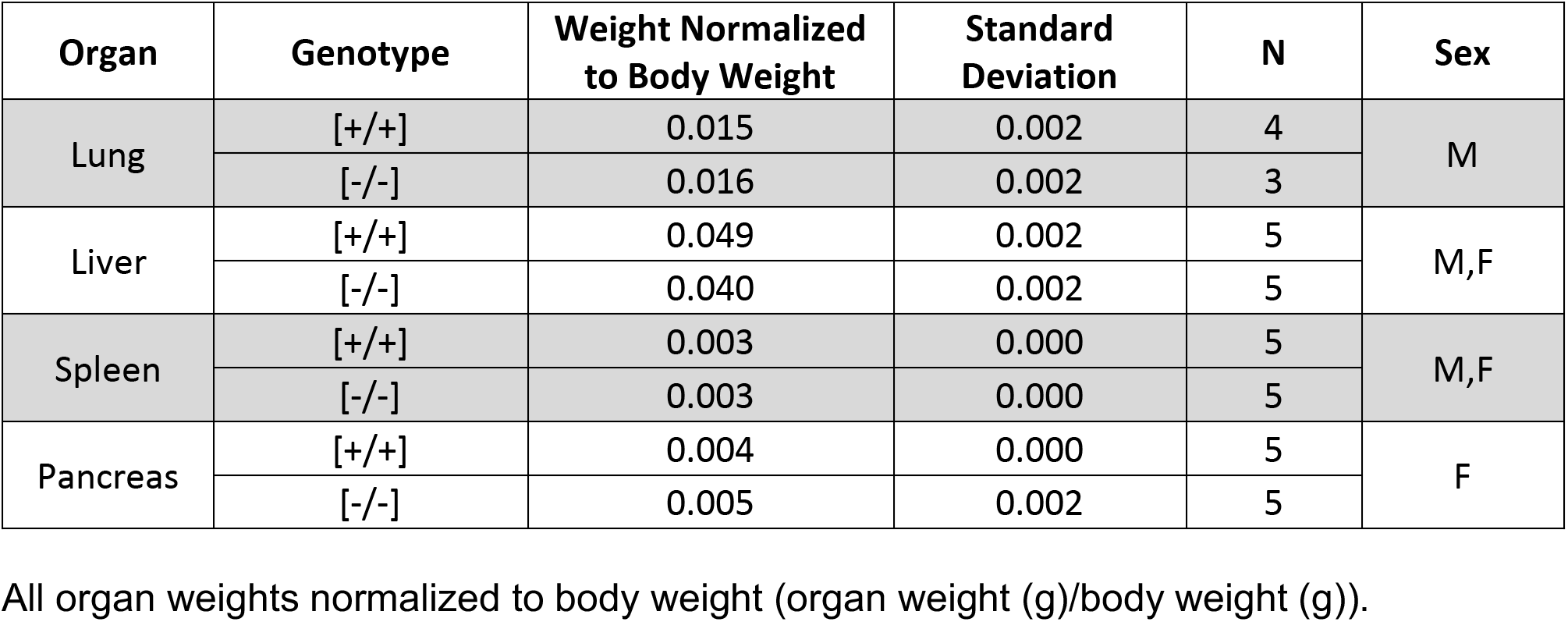
No significant impacts of *Zfp30* knockout observed on lung, liver, spleen, or pancreas weights.

**Supplementary Table 3:**
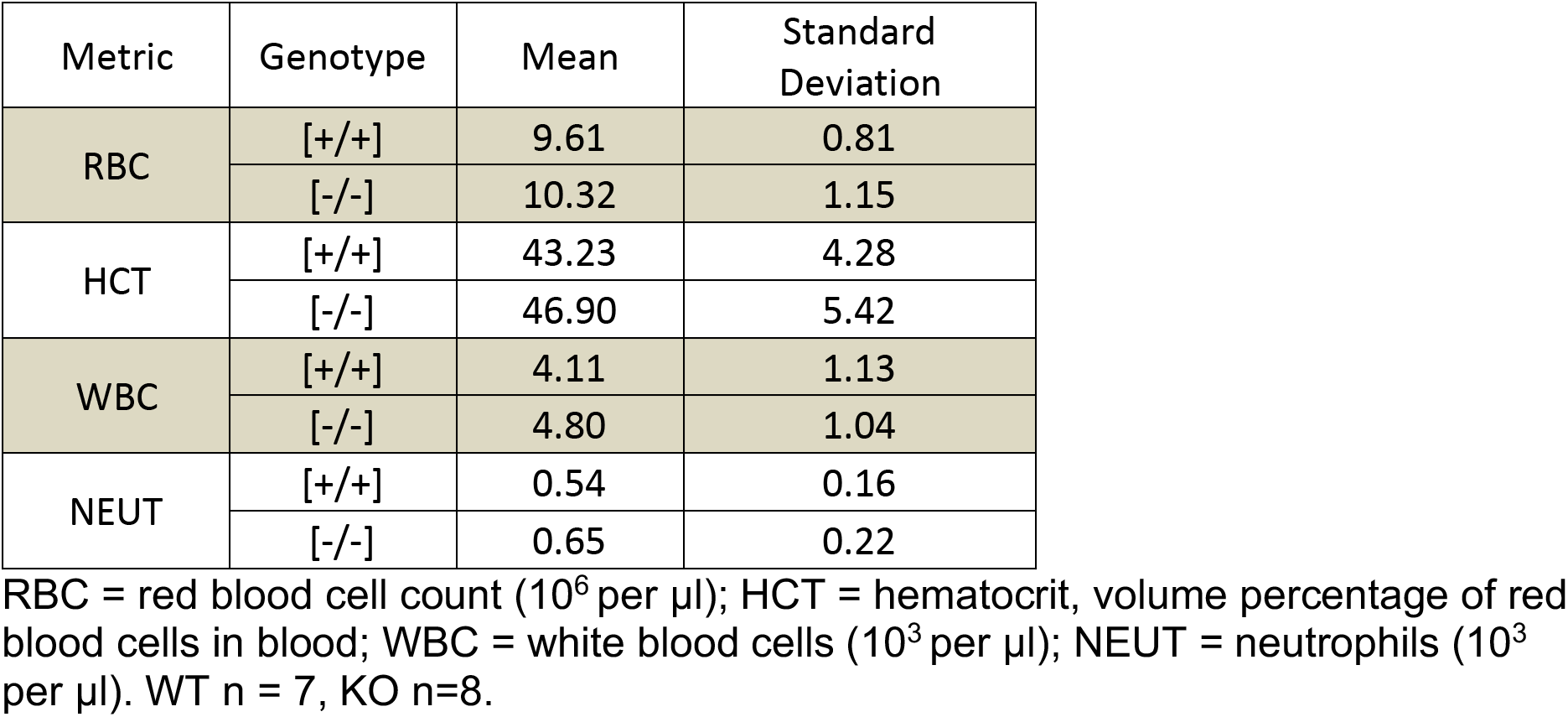
No significant impacts of *Zfp30* knockout observed on Complete Blood Count phenotypes.

**Supplementary Fig. 1.**
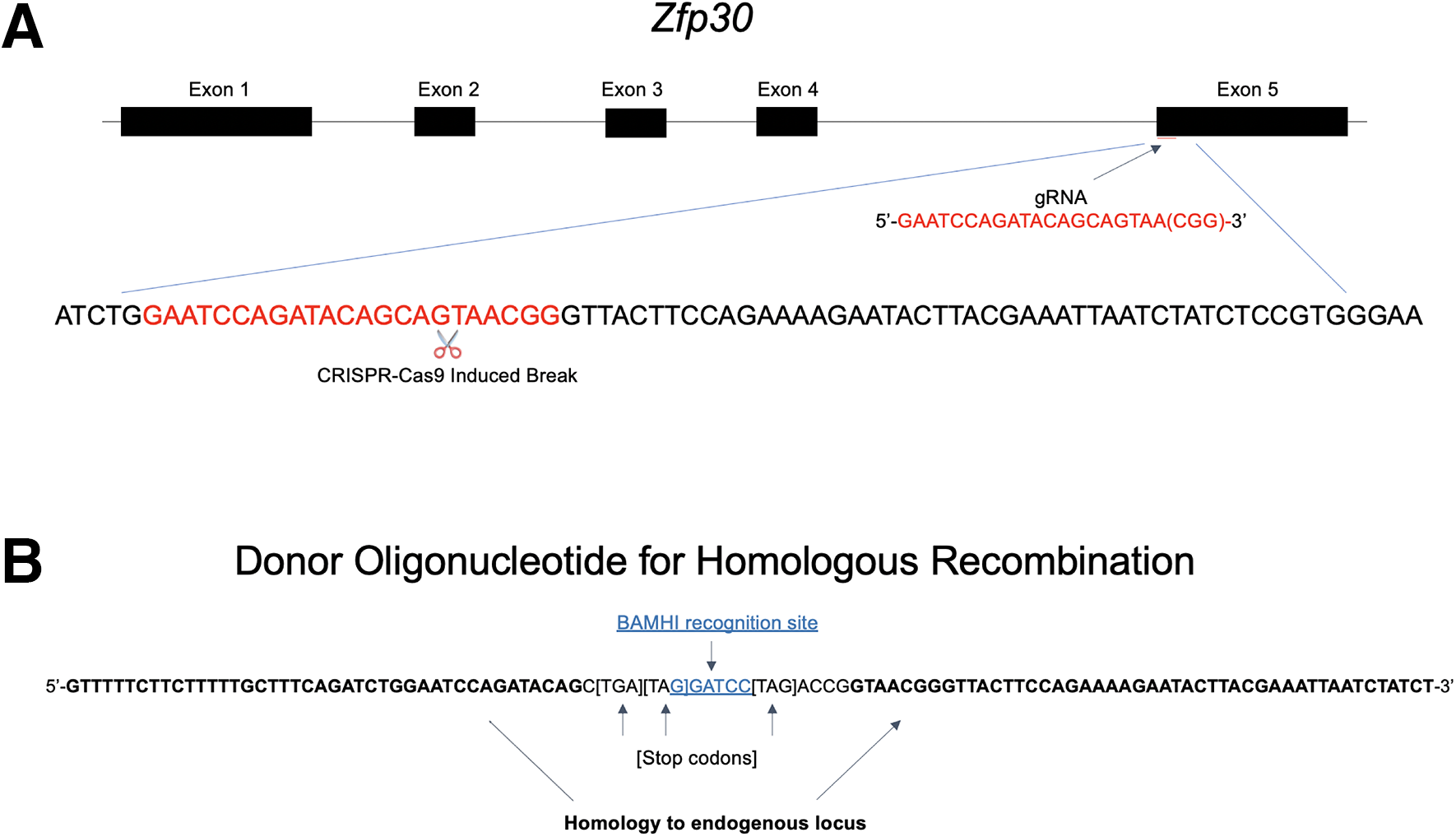
*Zfp30* CRISPR Strategy. A guide RNA was chosen to target exon 5 of *Zfp30* for breakage (A), and a donor oligonucleotide was designed to insert multiple stop codons and a BamHI target sequence at this site (B). Guide RNA sequence is shown in red; BamHI target sequence is shown in blue and underlined; stop codons are shown in brackets; homologous arms of donor oligo are shown in bold font.

**Supplementary Fig. 2.**
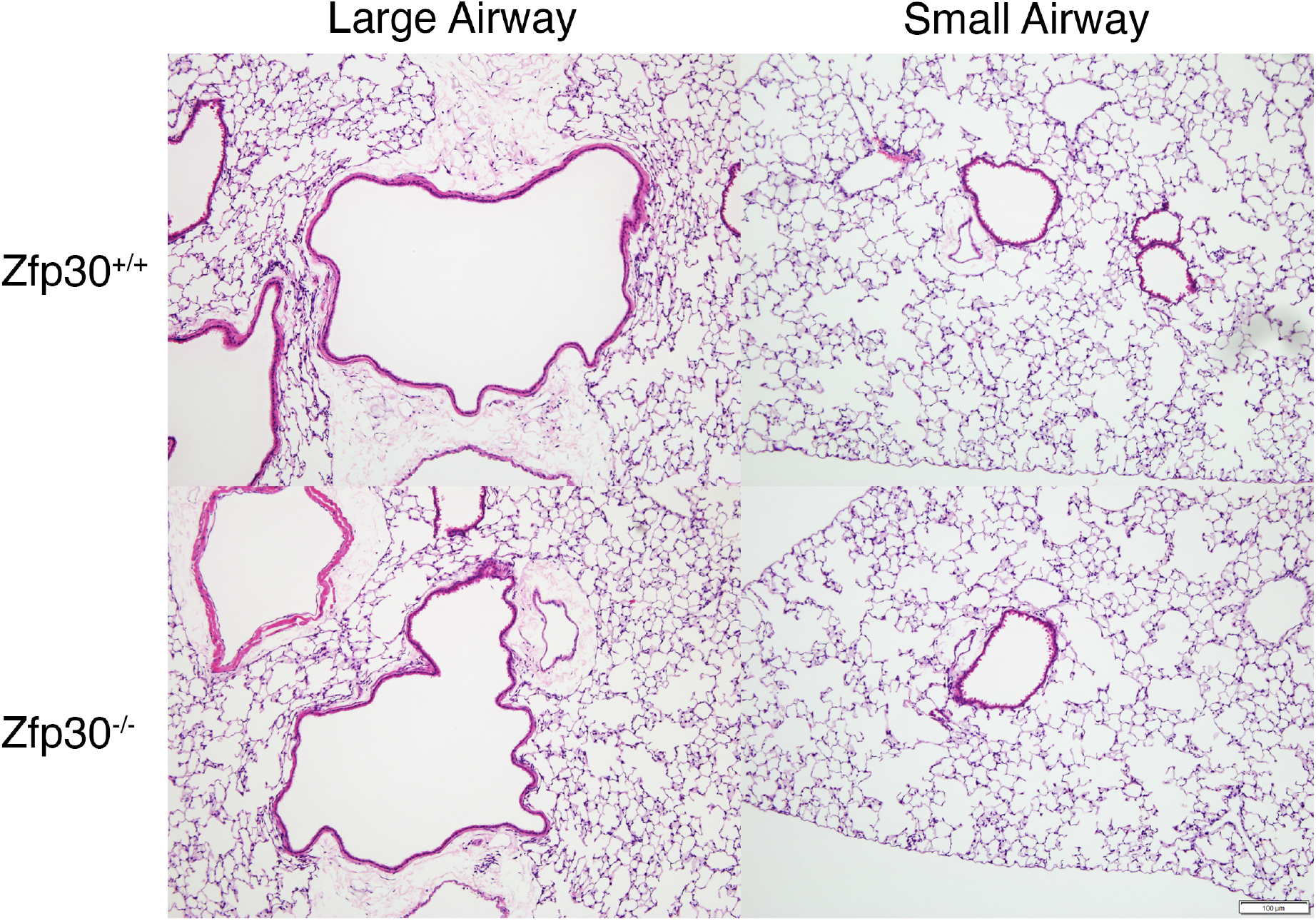
Histological analysis of lungs reveals no striking phenotypic differences by Zfp30 genotype. H&E staining was carried out on lung sections from *Zfp30*^+/+^ and *Zfp30*^-/-^ mice. No significant differences are seen in large airways (top left, bottom left) or among small airways and alveoli (top right, bottom right). Bar, 100μM.

